# A New Approach for Active Coronavirus Infection Identification by Targeting the Negative RNA Strand- A Replacement for the Current Positive RNA-based qPCR Detection Method

**DOI:** 10.1101/2023.01.22.525117

**Authors:** Darnell Davis, Hemayet Ullah

**Affiliations:** Department of Biology, Howard University, Washington, DC

## Abstract

This manuscript describes the development of an alternative method to detect active coronavirus infection, in light of the current COVID-19 pandemic caused by the SARS-CoV-2 virus. The pandemic, which was first identified in Wuhan, China in December 2019, has had a significant impact on global health as well as on the economy and daily life in the world. The current positive RNA-based detection systems are unable to discriminate between replicating and non-replicating viruses, complicating decisions related to quarantine and therapeutic interventions. The proposed method targets the negative strand of the virus and has the potential to effectively distinguish between active and inactive infections, which could provide a more accurate means of determining the spread of the virus and guide more effective public health measures during the current pandemic.

## Introduction

The COVID-19 pandemic has been exacerbated by the rapid spread of SARS-CoV-2 variants and increased transmission of infection due to the delay in the molecular detection of the infectious virus[1]. Though universal vaccination against the virus has occurred in some parts of the world, apprehensions of so called immune-escape-variants raise concern for a bleak future for the pandemic. The potential consequences of emerging variants are increased transmissibility, increased pathogenicity and the ability to escape natural or vaccine-induced immunity [2]. Several drugs with efficacy against the virus have been developed that are effective in reducing mortality and hospitalization. However, limited availability of drugs, contraindications, toxicity, rebound, etc. limit the widespread use of antiviral drugs [3, 4]. In addition, drug-escape-variants, like immune-escape-variants, can render existing treatments ineffective. All these issues make the effective management for the containment of the pandemic daunting and the impact of the continued pandemic is causing widespread disruption to daily life around the world.

In addition, recent reports of long-term viral persistence in multiple organs of COVID-infected persons are making containment efforts more complicated. For example, SARS-CoV-2 genome was found in the tonsils of 20% of 48 children that had tonsillectomies. None of the children had experienced signs or symptoms of COVID prior to the surgery [5]. The use of RT-PCR, immunohistochemistry (IHC), flow cytometry and neutralization assays on adenotonsillar tissues enabled detection of the SARS-CoV-2 genome [5]. Similarly, SARS-CoV-2 nucleocapsid protein specific immune-histological stain was seen in patients who underwent gastric and gallbladder surgery [6]. The presence of the virus protein was seen in patients who were COVID-19-free during surgery but were previously infected by the virus [6]. In addition, autopsies of several patients that died of COVID detected persistent SARS-CoV-2 RNA throughout the central nervous system as late as 230 days following symptom onset [7]. In another study, biopsies of patients with myocarditis showed the presence of SARS-CoV-2 spike protein and nucleocapsid antigens 1-5 months after infection. The maximal time after COVID-19 infection, when the virus was detected in the myocardium was 18 months [8].

Though the virus may be detected in different organs/tissues long after infection is over, whether these viruses represent replication-competent virus remains an urgent matter to resolve. Containment efforts to manage the pandemic will greatly be affected if infectious virions reside inside with patients long after recovering from COVID-19 infection. There are some claims about the presence of replication-competent virus by the detection of the sub-genomic positive mRNAs in the autopsied tissues [7]. In addition, isolation of replication-competent viruses in cell culture also potentially indicates the presence of replicating virus [7]. On the other hand, many investigators attribute the long-term presence of the SARS-CoV-2 to non-replicating viral fragments [8]. One study found spike particles trapped inside non-apoptosing non-classical monocytes of patients at up to 15 months post-infection [9].

In addition, current CDC guidelines requires 5-day isolation for individuals that test positive for COVID-19; however, many of these individuals have been found to remain positive on real time PCR tests for weeks after isolation, despite testing negative on antigen detection tests. Whether the persons that test positive on PCR remain infectious long after ending isolation remains a contentious issue. As such, it is now imperative to develop detection methods to distinguish replicating from non-replicating virus. Availability of such detection system would allow myriads of advantages, ranging from better informed decisions regarding quarantine policies to effective therapeutic decisions.

Realizing the acute need for such a detection system, here we propose and show the effectiveness of a detection system that can distinguish replicating virus from non-replicating virus. In this method, we show that the replicating mouse coronavirus continues to produce both their genome (positive RNA) and antigenome (negative RNA genome) throughout the infection process in cells. However, they cease to produce the antigenome when no opportunity to replicate in the cultured cells is present as cells get depleted due to the virus cytopathic effect. We could still detect the positive strand of the viral RNA in the live cell-depleted cell culture media, where no replication of the virus could take place due to the lack of host cells. Thus, the presence of the negative RNA strand may serve as a promising marker for viral replication competency and thus infectiousness. Here, we propose a negative RNA strand-specific detection method to replace or accompany current positive-strand specific real time PCR tests, in order to readily discern infectious disease through a simple detection test.

## Results

We propose that the coronavirus-negative mRNA strand can be established as a marker for replicating virus, which will aid in distinguishing replicating viruses from non-replicating viruses. To detect the negative strand, we used murine hepatitis virus, mouse coronavirus (MHV-A59) and used mouse fibroblast 17CL-1 cells to infect with the virus. We designed a specific primer to target the negative strand, such that the primer could be used to synthesize negative strand-specific cDNA. Similarly, we designed another primer to target the positive RNA strand and synthesize positive RNA strand-based cDNA. The negative cDNA primer sequence named NegcDNA.F spanned 30691 to 30713 bp of the complete genome of the murine hepatitis virus (accession #AY910861) with the following sequence:5’ TGAACCCACCAAAGATGTGTATG 3’. For the positive RNA strand, the primer (PoscDNA.R) sequence spanned the 31251 to 31273 bp of the complete genome of the murine hepatitis virus (accession #AY910861) with the following sequence: 5’ ACCCTGATGTGAGCTCTTCCCAG 3’ We developed a flow chart, as depicted in Figure 1, to detect the presence of the negative or positive strand in the MHV-A59 infected 17CL-1 cell lines.

**Figure 1:**
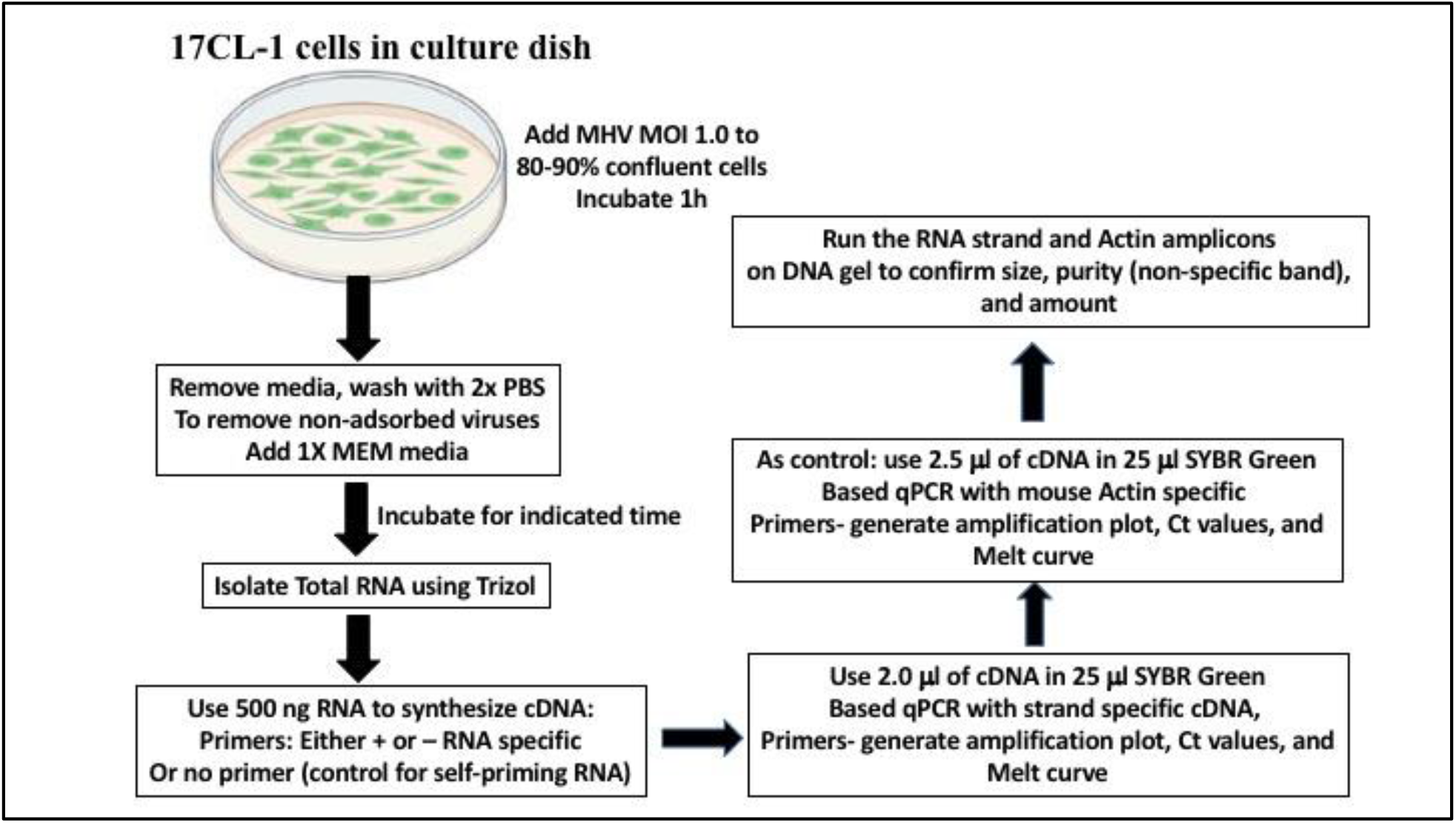
Flow chart depicting the experimental steps for detecting the mouse coronavirus MHV-A59 positive or negative RNA strand. Self-priming RNA based cDNA developed without any primer shows the background level of expression. RT-PCR refers to the Reverse Transcriptase based PCR where is real time PCR is described as qPCR.

MHV have been widely used as a model for the family of enveloped plus RNA viruses from Coronaviridae, and the MHV-A59 strain is the prototype coronavirus (CoV). We used strain MHV-A59 harboring an eGFP fluorescent tag inserted by replacing a pseudogene-ORF4 [10]. The virus was obtained from NIAID BEI resources, and a stock was prepared in the laboratory by infecting the mouse fibroblast cell line 17CL-1.

Following the flow chart, we used 500 ng of the total RNA isolated after two hours of infection by 1.0 MOI MHV-A59 in 17CL-1 cells to synthesize the strand-specific cDNA. Using two primers, NegcDNA.F and PoscDNA.R, we amplified an approximately 600 bp long PCR product from the respective cDNA. As a control for the loading, we used mouse actin primers named mActin.F with the sequence 5′-GGCTGTATTCCCCTCCATCG-3′ and mActin.R with the sequence 5′-CCAGTTGGTAACAATGCCATGT-3′ to amplify a 154 bp long PCR product. Both PCR products are depicted in Fig. 2, which shows that we could easily detect both the negative and the positive strand-specific bands from the cDNA synthesized two hours following viral infection.

**Figure 2:**
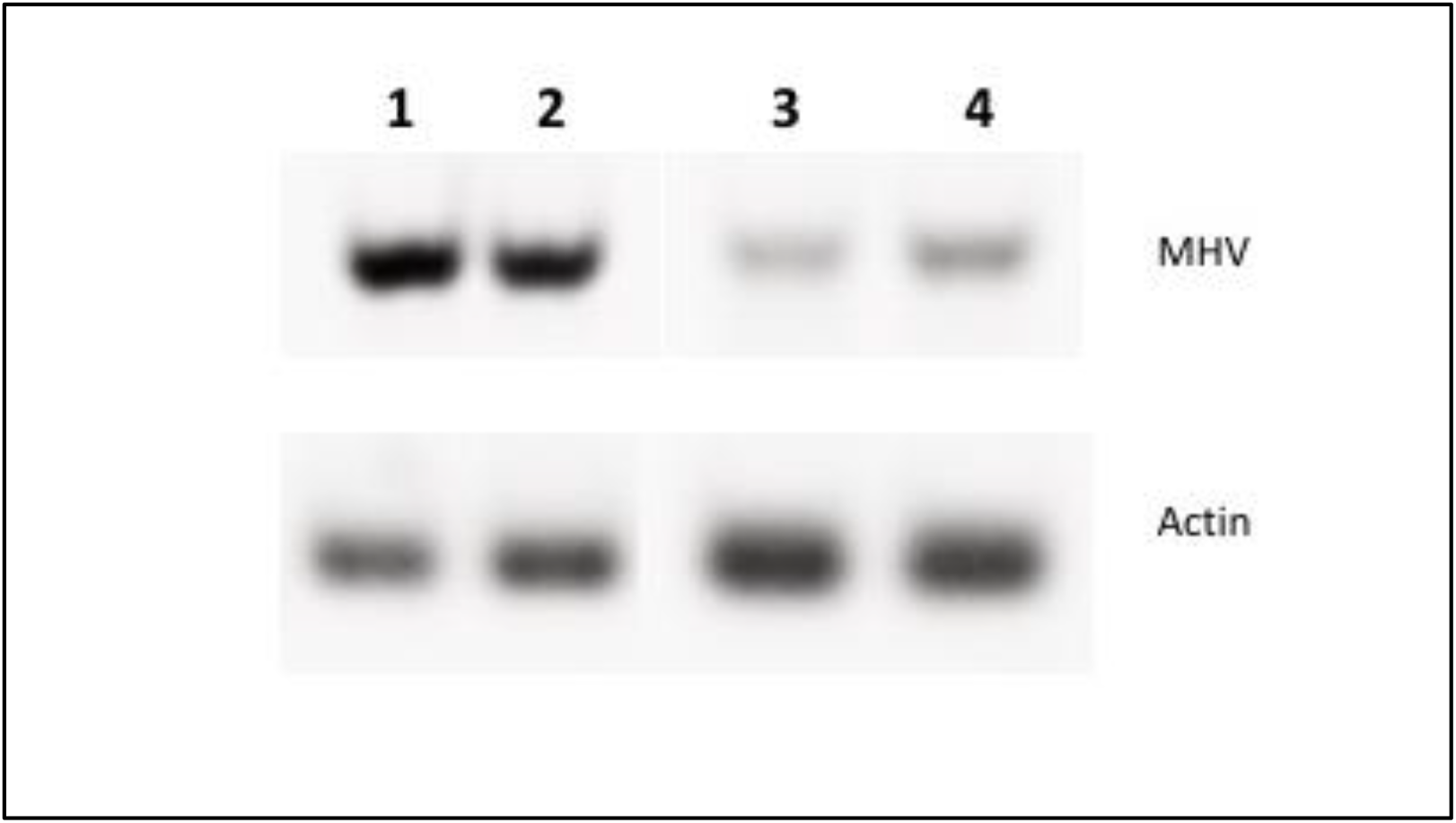
RT-PCR of the CR product specific to the negative RNA strand (lane 1 and 2 in duplicate) and the positive RNA strand band after 2h of infection of the 17CL-1 cells with the MOI 1.0 of mouse coronavirus- MHV-A59. The lower panel shows the loading control by amplifying the mouse actin band of 154 bp length.

Although the intensity of the negative strand-specific strand indicated more negative strand synthesis immediately after infection, more studies are needed to find any biological implications for this expression. However, as our detection system was able to detect the negative strand-specific band more easily, it can be safely inferred that this system can be robustly used to detect the negative strands of a coronavirus.

Because antigenome production is necessary to sustain the replication of the virus, we tested whether we could detect the negative strand continuously when active infection was underway. As active infection requires the presence of host cells and the virus cytopathic effect depletes the host cells over time, we isolated RNA samples from different time points of infection, as well as when almost all host cells were depleted from the cytopathic effect of the infecting virus. Although we isolated from 2h until the end of the total cytopathic effect, Fig. 3 shows some of the stages of the cells from which total RNA was isolated for strand-specific RNA identification.

**Figure 3:**
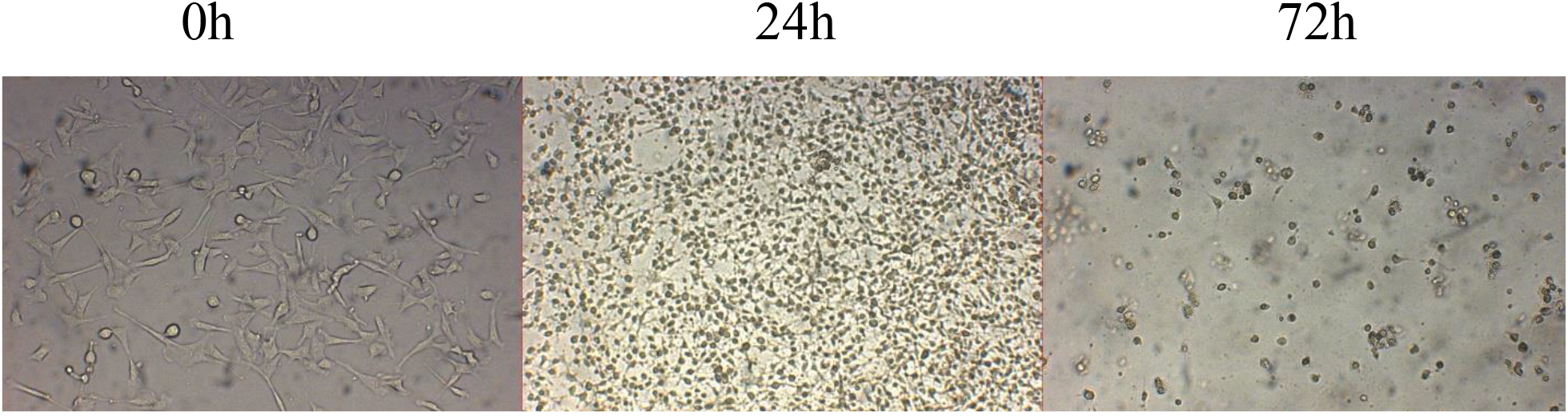
Developmental stages of the host cells(17CL-1) post infection with the MHV-A59. Note the host cells shows extensive signs of infection within the 24h post infection and by the time of 72h post infection, hardly any host cells can be seen. The Cytopathic effect is completed by 72h post infection.

Fig. 3 shows that after 3 days of infection with an MOI 1.0 MHV-A59 virus, almost all the host cells were depleted. In all samples throughout the infection, we detected both positive and negative RNA strands. However, we tested whether both strands could be detected in RNA samples isolated from host cell-depleted media collected after 72h post infection. As no host cells were available, no active infection demonstrably occurred in those samples. Here, we also use no primer cDNA synthesis from the same RNA sample as a control for the widely reported ‘self-priming’ based cDNA synthesis [11]. Fig. 4 shows the RT-PCR-based detection of

**Figure 4:**
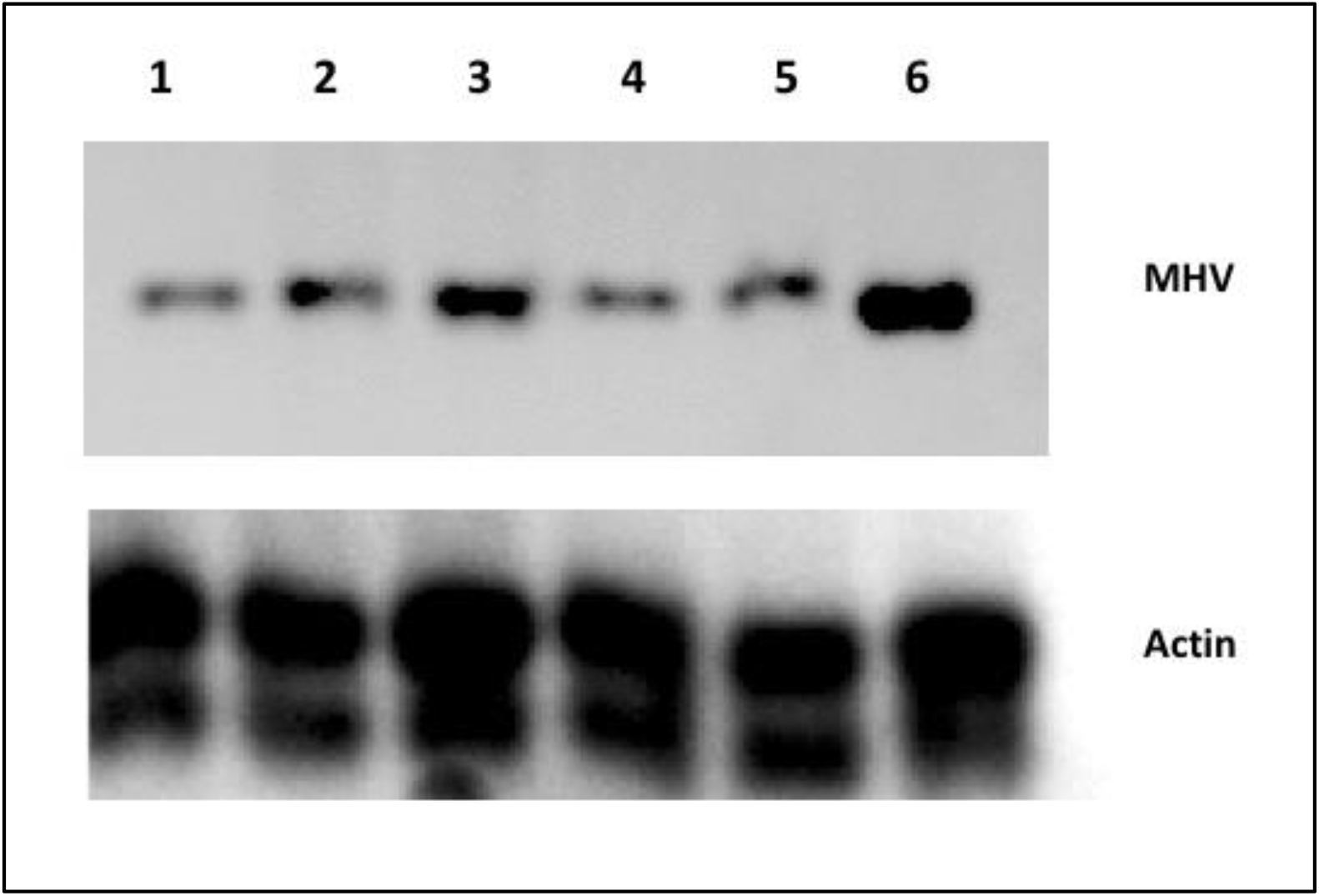
RT-PCR based PCR product targeting the positive or the negative RNA strand. Lane 1 and 4-24h and 72h post infection with self-priming cDNA samples respectively. Lane 2 and 5-24h and 72h post infection with negative RNA specific cDNA samples, and lane 3 and 6-24h and 72h post infection with positive RNA strand specific cDNA samples. Lower panel shows mouse actin (154 bp) from the same RNA isolated after 24h and 72h post infection samples.

MHV-A59 in the no-primer, negative, and positive cDNA from the 72h post infection media. Similar to the samples in Fig 1, PCR amplification was carried out using the same two primers, NegcDNA.F and PoscDNA.R, which amplify a 600 bp product. Lanes 1, 2, and 3 represent cDNA synthesized 24h post infection while lane 4, 5, and 5 represent cDNA synthesized 72 h post-infection. Lanes 1 and 4 represent the band detected from the no-primer cDNA, showing the background level of the PCR band amplified from the self-primed cDNA. Lanes 2 and 5 represent the bands from the negative RNA-specific cDNA. Note that lanes 4 and 5 show almost identical levels of expression at 72h post infection-indicating that there was no amplification from the minus RNA strand, as the expression represents the background level of expression, similar to the no-primer expression. This establishes that no negative RNA strands can be detected when no host cells are available. On the contrary, an intense band was seen from the positive strand-specific cDNA at 72h post infection. As there were no host cells available (Fig. 4) at 72h post infection, this band likely originates from the broken-down genome, remnant/residual virus fragments, or non-replicating virions in the media. Thus, positive RNA can be detected from non-replicating viruses, but to detect the negative strand, a sample from replicating viruses within host cells is needed. As a control, we used mouse actin primers for the same cDNA samples (lower panel).

To confirm the RT-PCR results, we used the same cDNA (72h post-infection) in the qPCR experiment, and the results are depicted in Fig. 5. In concordance with the RT-PCR experiment, serial dilution of the cDNA produced the exact same level of Cycle Threshold (Ct) values as that in no-primer and negative strand specific-cDNA conditions, while the positive RNA-specific cDNA sample showed lower Ct values, indicating higher expression. Here, we used a higher level of cDNA dilutions (0.01 and 0.1 dilution) to obtain robust amplification. Instead of actin, we quantified the three cDNA samples in a nanodrop machine, and the same level of concentrations in each of the cDNA samples is indicated in the nanodrop light absorption chart (Fig. 6B). The melting curve for all samples clearly shows that all samples amplified the same targeted band without amplifying any non-specific bands.

**Figure 5:**
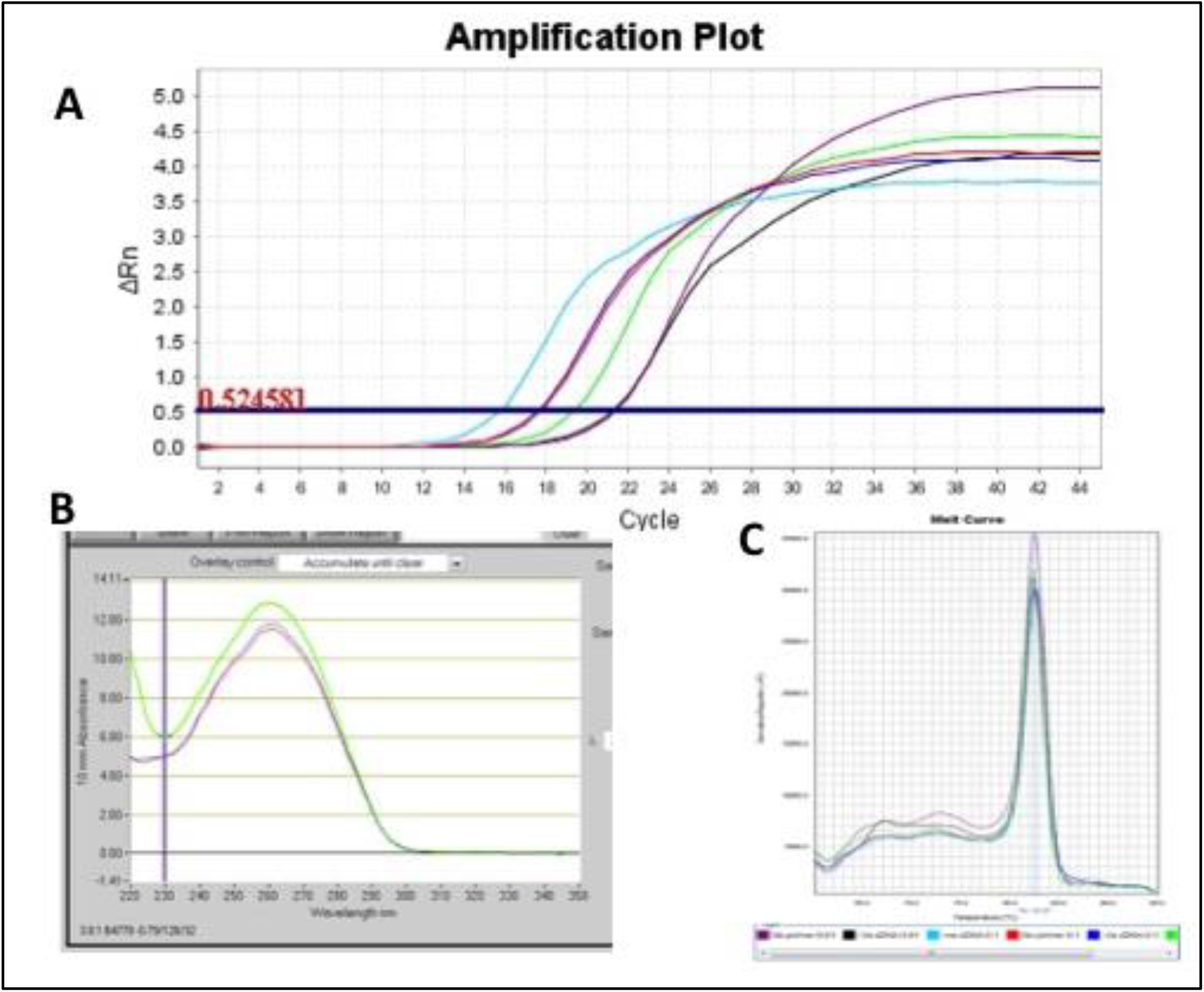
qPCR amplification plot of the negative and positive RNA strand specific cDNA samples. cDNAs from the 72h post infection samples were shown which was amplified at 0.01 and 0.1 dilutions of the cDNAs. Purple and black amplification plots in panel A represent 0.01 dilution samples from self-priming and negative-RNA strand specific cDNAs while green amplification plot represents the positive RNA strand specific cDNA sample. Red and blue amplification plots represent the self-priming and negative strand specific cDNA samples while the turquoise amplification plot represent the positive RNA strand specific cDNA sample at 0.1 dilution. Panel B shows the absorption spectra of all three cDNA samples. Panel C shows the melting curve of the amplified products in the panel A.

Thus, we show here both by RT-PCR and qPCR that when the virus is not replicating actively, we can still detect virus-positive strand-specific signals, whereas only the background level of the signal (similar to the no-primer cDNA) can be detected from the negative strand-specific RNA.

## Discussion

The current pandemic has had a profound impact on the world, with the virus spreading rapidly and causing significant illness and death since it started in 2020. To counter the pandemic, researchers and healthcare professionals have been developing new tools and technologies for the detection and diagnosis of the virus. However, the inability to distinguish replicating viruses from non-replicating viruses limits the utility of the current detection methods. We describe here the development of an alternative active coronavirus detection system that uses the negative mRNA strand as a marker for the presence of a replicating virus. This approach offers several potential advantages over existing methods, which will aid in the effective management of the current pandemic situation.

One of the key advantages of using the negative mRNA strand as a target for detection is its ability to distinguish between active and non-active infections. By targeting the negative strand, this detection approach should be able to detect only those individuals who are currently infected with actively replicating virus, and thus are capable of transmitting it to others. This is an important distinction, as individuals who have recovered from the virus may still test positive using other detection methods targeting the positive strand, but that may not necessarily indicate infectiousness, as remnant/residual virus genome fragments can yield positive results in the absence of any replication.

Furthermore, the presence of a negative strand is a clear indicator of active infection. If an individual’s sample does not contain the negative strand, this indicates that they are not currently infected with any actively replicating virus and do not need to be quarantined. This can help reduce the burden on healthcare systems and prevent the unnecessary isolation of individuals.

Another key strength of this approach is its high sensitivity and specificity. By targeting the negative mRNA strand, our system was able to accurately detect the presence of the virus, even at low levels. This is particularly important in the early stages of the disease, when the amount of virus in an individual sample may be low. Additionally, the use of the negative strand as a target allows for the specific detection of coronavirus, reducing the potential for false positives.

Additionally, the use of the negative strand of coronavirus RNA as a marker for replicating viruses may be useful for monitoring the effectiveness of antiviral therapies. By measuring changes in the levels of negative-strand RNA over time, clinicians can determine whether a particular treatment effectively reduces the number of replicating viruses in the body. This information can be used to guide treatment decisions and optimize patient outcomes.

Development of this detection approach for active infection can have wide applications as a very large group of viruses belonging to many families are found to replicate through positive to negative RNA strand method. Along with coronaviruses, other positive RNA viruses that cause epidemic diseases in humans include encephalitis, hepatitis, polyarthritis, yellow fever, dengue fever, poliomyelitis, and common cold-causing viruses. Widespread use of the negative strand-specific detection system will not only address the most pressing issues of the current pandemic, but will also present opportunities to address other disease-causing RNA viruses as well.

The use of the negative mRNA strand as a marker allows for the direct detection of replicating viruses, rather than relying on indirect indicators, such as the presence of viral proteins or antigens. This could result in a more sensitive and specific detection method. The use of the negative mRNA strand as a marker allows for real-time monitoring of virus replication, as the negative strand is produced during active viral replication. This could provide valuable information for disease management and control.

Overall, our findings support the establishment of the negative mRNA strand as a valuable target for active coronavirus detection. Previously, we showed how the negative strand’s unique 5’ polyU tag can be used to inhibit coronavirus replication without allowing any mutation-based variant development [12]. Further research is needed to evaluate the performance of this system in larger, more diverse populations, and to assess its potential for use in real-world settings. However, our results suggest that this approach has the potential to significantly improve prospects for the rapid identification of infected individuals and ultimately, in combatting spread of the virus.

## Acknowledgement

The following reagents were obtained from BEI Resources (NIAID/NIH): Recombinant Murine Coronavirus MHV-A59 with Enhanced Green Fluorescent Protein (eGFP) (Cat# NR-53716), Murine 17Cl-1 Cell Line (derived from 3T3 cells) (Cat# NR-53719). No funding was used to develop the findings of this project.

## Notes

### Competing Interest Statement

The authors have declared no competing interest.

## References

1. Frampton, D.; Rampling, T.; Cross, A.; Bailey, H.; Heaney, J.; Byott, M.; Scott, R.; Sconza, R.; Price, J.; Margaritis, M.; Bergstrom, M.; Spyer, M. J.; Miralhes, P. B.; Grant, P.; Kirk, S.; Valerio, C.; Mangera, Z.; Prabhahar, T.; Moreno-Cuesta, J.; Arulkumaran, N.; Singer, M.; Shin, G. Y.; Sanchez, E.; Paraskevopoulou, S. M.; Pillay, D.; McKendry, R. A.; Mirfenderesky, M.; Houlihan, C. F.; Nastouli, E., Genomic characteristics and clinical effect of the emergent SARS-CoV-2 B.1.1.7 lineage in London, UK: a whole-genome sequencing and hospital-based cohort study. Lancet Infect Dis 2021.

2. Vincent Garcia, V. V., Laurent Peillard, Alaa Ramdani, Sofiane Mohamed, Philippe Halfon, First description of two immune escape indian B.1.1.420 and B.1.617.1 SARS-CoV2 variants in France. bioRxiv 2021, 2021.05.12.443357.

3. Wen, W.; Chen, C.; Tang, J.; Wang, C.; Zhou, M.; Cheng, Y.; Zhou, X.; Wu, Q.; Zhang, X.; Feng, Z.; Wang, M.; Mao, Q., Efficacy and safety of three new oral antiviral treatment (molnupiravir, fluvoxamine and Paxlovid) for COVID-19:a meta-analysis. Ann Med 2022, 54, (1), 516–523.

4. Wang, L.; Berger, N. A.; Davis, P. B.; Kaelber, D. C.; Volkow, N. D.; Xu, R., COVID-19 rebound after Paxlovid and Molnupiravir during January-June 2022. medRxiv 2022.

5. Miura CS, Lima TM, Martins RB, Jorge DMM, Tamashiro E, Anselmo-Lima WT, Arruda E, Valera FCP. Asymptomatic SARS-COV-2 infection in children’s tonsils. Braz J Otorhinolaryngol. 2022 November-December;88:9. doi: 10.1016/j.bjorl.2022.10.016. Epub 2022 Oct 20. PMCID: PMC9582977.

6. Hany, M.; Zidan, A.; Gaballa, M.; Ibrahim, M.; Agayby, A. S. S.; Abouelnasr, A. A.; Sheta, E.; Torensma, B., Lingering SARS-CoV-2 in Gastric and Gallbladder Tissues of Patients with Previous COVID-19 Infection Undergoing Bariatric Surgery. Obes Surg 2023, 33, (1), 139–148.

7. Stein, S. R.; Ramelli, S. C.; Grazioli, A.; Chung, J. Y.; Singh, M.; Yinda, C. K.; Winkler, C. W.; Sun, J.; Dickey, J. M.; Ylaya, K.; Ko, S. H.; Platt, A. P.; Burbelo, P. D.; Quezado, M.; Pittaluga, S.; Purcell, M.; Munster, V. J.; Belinky, F.; Ramos-Benitez, M. J.; Boritz, E. A.; Lach, I. A.; Herr, D. L.; Rabin, J.; Saharia, K. K.; Madathil, R. J.; Tabatabai, A.; Soherwardi, S.; McCurdy, M. T.; Consortium, N. C.-A.; Peterson, K. E.; Cohen, J. I.; de Wit, E.; Vannella, K. M.; Hewitt, S. M.; Kleiner, D. E.; Chertow, D. S., SARS-CoV-2 infection and persistence in the human body and brain at autopsy. Nature 2022, 612, (7941), 758–763.

8. Blagova, O.; Lutokhina, Y.; Kogan, E.; Kukleva, A.; Ainetdinova, D.; Novosadov, V.; Rud, R.; Savina, P.; Zaitsev, A.; Fomin, V., Chronic biopsy proven post-COVID myoendocarditis with SARS-Cov-2 persistence and high level of antiheart antibodies. Clin Cardiol 2022, 45, (9), 952–959.

9. Patterson, B. K.; Francisco, E. B.; Yogendra, R.; Long, E.; Pise, A.; Rodrigues, H.; Hall, E.; Herrera, M.; Parikh, P.; Guevara-Coto, J.; Triche, T. J.; Scott, P.; Hekmati, S.; Maglinte, D.; Chang, X.; Mora-Rodriguez, R. A.; Mora, J., Persistence of SARS CoV-2 S1 Protein in CD16+ Monocytes in Post-Acute Sequelae of COVID-19 (PASC) up to 15 Months Post-Infection. Front Immunol 2021, 12, 746021.

10. Das Sarma, J.; Scheen, E.; Seo, S. H.; Koval, M.; Weiss, S. R., Enhanced green fluorescent protein expression may be used to monitor murine coronavirus spread in vitro and in the mouse central nervous system. J Neurovirol 2002, 8, (5), 381–91.

11. Zhang, C.; Wu, H. N.; Zhang, Y. Q.; Shen, J. G.; Li, W. M., Self-priming on the plant viral RNAs during reverse transcription-PCR. Acta Virol 2015, 59, (1), 92–7.

12. Ullah, H.; Averick, S.; Tang, Q., Crippled Coronavirus: 5’-PolyU targeted Oligo prevents development of infectious Virions. Biorxiv 2021, 2022.03.04.483076, (doi: https://doi.org/10.1101/2022.03.04.483076).

